# Relative Effects of Habitat Amount and Fragmentation *Per Se* on the Genetic Diversity of The Glanville Fritillary Butterfly

**DOI:** 10.1101/2024.11.06.622224

**Authors:** Lola Fernandez Multigner, Audrey Bras, Michelle DiLeo, Marjo Saastamoinen

## Abstract

Habitat loss and fragmentation are considered the key drivers of biodiversity loss. Understanding their relative roles is difficult as habitat loss and fragmentation tend to co-occur. It has been proposed that the total habitat amount available in the local landscape mainly drives species richness while fragmentation *per se* – the breaking apart of habitat independent of habitat amount - has negligible or even a positive effect on biodiversity. Several studies support this at the species richness level. Yet, the potential effects of fragmentation *per se* on genetic diversity at the landscape scale are understudied. Using the Glanville fritillary butterfly metapopulation in the Åland islands, we tested the effects of fragmentation *per se* on genetic diversity using a landscape-based approach and 2,610 individuals genotyped at 40 neutral SNP markers. We assessed the independent effect of habitat amount and fragmentation (i.e. number of patches) within the local landscape on the focal patch genetic diversity. The amount of habitat in the local landscape had a positive effect on genetic diversity, while fragmentation *per se* had a more negligible impact on the genetic diversity. Our results thus highlight that all fragments, even the small ones, likely contribute to the maintenance of genetic diversity of the focal population.

## Introduction

Biodiversity loss is among the greatest challenges human societies are currently facing. Predicting the magnitude and drivers affecting biodiversity loss is urgently needed to prioritize conservation efforts. However, biodiversity loss is often measured as the loss of species diversity, with indicators such as genetic diversity often being entirely neglected (Pereira et al. 2013). Genetic diversity is directly linked with evolutionary potential of a species and its adaptability (Hu et al. 2021), hence genetic diversity loss is expected to affect a population’s or species’ long term survival. It has recently been estimated that more than 10% of genetic diversity might have already been lost for many species due to human activities such as habitat destruction (Exposito-Alonso et al. 2022).

Habitat loss and fragmentation are often cited together as main drivers of biodiversity loss (Tilman et al. 1994; Huxel & Hastings 1999). Based on their current increase rates, both factors are expected to continue causing most of the biodiversity loss in the future (Visconti et al. 2016). However, habitat loss and fragmentation are two distinct processes with potentially varying impacts on populations (Fahrig 2003, 2017). Therefore, disentangling the threats each process imposes is needed for effectively mitigating biodiversity loss. Habitat loss refers to the reduction of habitat amount within a landscape (Martin et al. 2021), and can negatively affect various hierarchical levels of biodiversity, such as species richness (i.e. largely due to the general species-area relationship; Lomolino 2000), or intraspecific genetic diversity (Exposito-Alonso et al. 2022). Habitat fragmentation, on the other hand, has widely been used as an umbrella term for many ecological processes accompanying alteration of landscapes by humans (Lindenmayer & Fischer 2007). Its different definitions and effects on biodiversity have been a continuous debate (see for instance: Fahrig 2003, 2013; Valente et al. 2023). However, it is usually defined as a previously continuous habitat being destroyed into smaller unconnected habitats. As habitat fragmentation usually happens together with habitat loss, assessments of its impacts alone on biodiversity are difficult (Fahrig 2003; Ewers & Didham 2006).

This general coupling of habitat loss and habitat fragmentation has challenged the general assumption of the solely negative effect of habitat fragmentation on biodiversity. It has been suggested that the configuration of habitat in the landscape generally has little or no effect on species richness in sampled sites, provided that the habitat amount in the landscape stays constant (Fahrig 2003, 2013, 2017). In this context, habitat fragmentation, or fragmentation *per se*, is independent of the effects of habitat amount, and the fragmentation process does not necessarily imply habitat loss (Martin et al. 2021). Studies assessing the effects of fragmentation *per se* on biodiversity have principally focused on species richness changes depending on the amount of habitat or degree of fragmentation in a landscape (Haddad et al. 2017; Seibold et al. 2017; Watling et al. 2020; Herrero-Jáuregui et al. 2022), while impacts on genetic diversity have been largely neglected (but see Jackson & Fahrig 2014; González-Fernández et al. 2019). Disentangling the effects of habitat amount and fragmentation *per se* on genetic diversity is, however, equally important to determine their respective direction and magnitude.

Landscape genetic studies have largely looked at the effects of landscape features on genetic diversity with a focus on habitat configuration, but the considered scale has predominately been at the patch level (DiLeo & Wagner 2016). Such patch-scale approaches cannot determine impacts of fragmentation *per se* on genetic diversity, as measures of the fragmentation of the habitat are confounded with habitat amount (Fahrig 2019). Determining potential effects of fragmentation *per se* over habitat amount on genetic diversity at the landscape level will allow the development of more informed conservation management strategies that consider species ability to adapt to changing environmental conditions (DiLeo & Wagner 2016; Savary et al. 2022). Indeed, the size of a fragment can limit the local population size it holds, which, when reduced, could cause a loss in genetic variation through genetic drift (Frankham 1996). Thus, a population subject to habitat reduction and coupled with habitat fragmentation can show lower genetic diversity with an increased genetic differentiation among the subpopulations (Keyghobadi et al. 2005). On the other hand, if the amount of habitat is equal within the landscape, and the remaining habitats are well enough connected to each other, populations living in more fragmented habitats could maintain an equal or have even a higher genetic diversity due to spatially varying selection (Hanski et al. 2011; Cheptou et al. 2017).

The Glanville fritillary butterfly (*Melitaea cinxia*) in the Finnish Åland Islands archipelago has become a model system to study the effect of fragmentation on metapopulation dynamics, and the ecology and genetics of the butterfly have been well studied (Ojanen et al. 2013; Hanski et al. 2017). Within this metapopulation, each suitable habitat across the landscape has been characterized (Hanski et al. 2017), and since 1993, the set of approximately 4,000 habitat patches has been annually systematically monitored for the presence of conspicuous larval families (Hanski et al. 1996; Ojanen et al. 2013). The effect of habitat area, connectivity, and configuration on the metapopulation dynamics (i.e. occupancy, abundance, local extinction, and re-colonization) have been extensively studied using a patch-based approach (Hanski et al. 1994, 1995; Saccheri et al. 1998; Hanski 2011b; Schulz et al. 2020; Bergen et al. 2020). The spatial configuration and habitat quality rather than the pooled habitat area have been shown to predict metapopulation size and persistence in combination with genetic variation in species’ dispersal capacity (Hanski et al. 2017; Fountain et al. 2018; DiLeo et al. 2018). A recent study using a landscape-based approach has also demonstrated the positive effect of fragmentation *per se* on patch occupancy and colonization (Galán-Acedo et al. 2024). Yet, little is known about the spatial variation of genetic diversity at a landscape scale, nor the effect of fragmentation *per se* on genetic diversity which influences the butterfly’s population performance (DiLeo et al. 2024).

The long-term monitoring survey together with spatio-temporal genetic data of the Glanville fritillary butterfly over the Åland islands offers a comprehensive dataset to assess how habitat amount and fragmentation *per se* are impacting a species’ genetic diversity (Hanski 2015). We used the extensive characterization of the butterfly’s habitat in the Åland archipelago (Ojanen et al. 2013), and an intensive sampling carried out during 2 consecutive years during which 4,862 larval family nests were sampled and 12,938 individuals were genotyped for 40 neutral SNP markers (DiLeo et al. 2018). Using a landscape-based approach, we first defined the scale of effect which corresponds to a buffer radius for which landscape variables best predict local population responses, following Martin et al. (2021). We then disentangled the effects of habitat amount from fragmentation *per se* on the neutral genetic diversity of the focal patch. As the amount of habitat available is likely to impact the genetic diversity greatly, we predicted higher levels of inbreeding with a decreasing amount of habitat available in the landscape. For a constant amount of habitat within the landscape, we further expected a neutral effect of fragmentation *per se* on the genetic diversity.

## Material and Methods

### Study system and landscape characterization

The Glanville fritillary butterfly is distributed across Europe up to the Åland archipelago in southern Finland. Within the Åland islands, the butterfly lives in a highly fragmented landscape and persists as a classic metapopulation (Hanski et al. 1994). The butterfly has one generation per year and overwinters as larvae in a winter nest. It has two larval host plants, *Plantago lanceolata* and *Veronica spicata*, growing on dry meadows and pastures, which often represent identifiable and discrete habitat patches (Nieminen et al. 2004).

In the Åland islands, each suitable habitat patch for *Melitaea cinxia* has been characterized and georeferenced based on the availability of one or both host plant species, corresponding to over 4,000 well-defined habitat patches (Ojanen et al. 2013). The habitat patches vary in size and are spread out within a heterogeneous landscape that includes agricultural lands, rocky areas, woodland clearings, and urbanized areas. Each potential habitat patch has been systematically visited every fall since 1993 to assess the presence of winter nests and their abundance. This information is used to evaluate the butterfly’s population demography, patch occupancy, patch extinctions and patch recolonizations (Nieminen et al. 2004). Using this extensive dataset, we classified the local landscapes based on the cartography of all these discrete habitat patches (Fig. 1).

**Figure 1.**
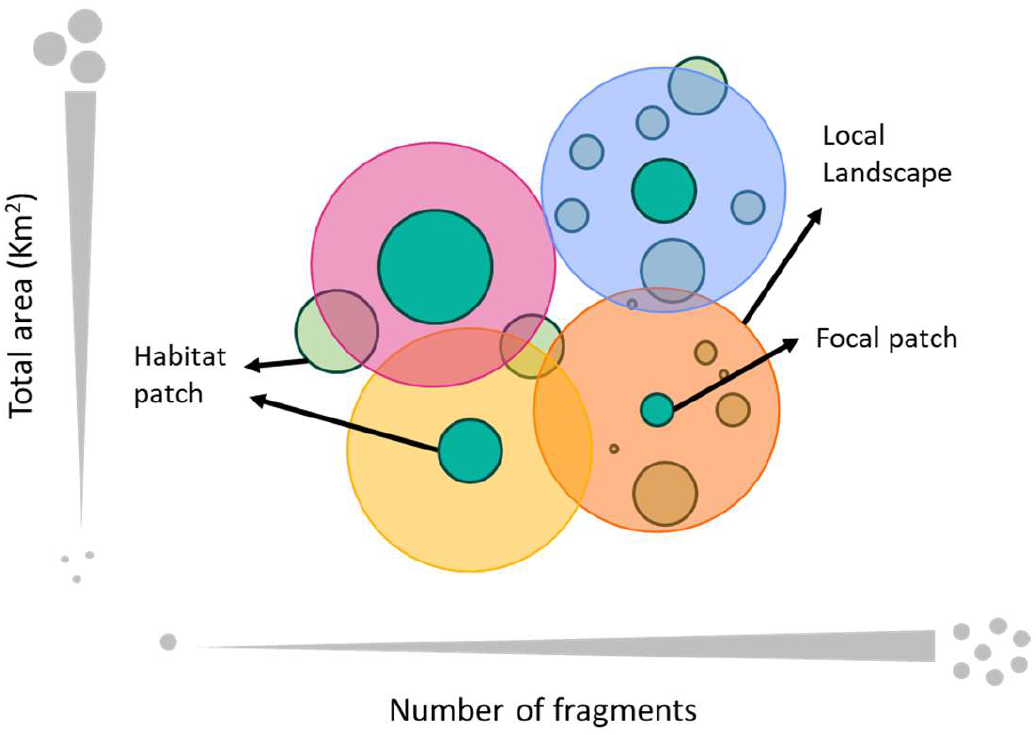
Schematic representation of the landscape categorization into habitat patches, focal patches and local landscapes with varying fragmentation and habitat amount.

### Genetic diversity indices

To compute genetic diversity indices, we utilized an existing dataset from years 2011 and 2012, when up to three larvae per nest across the Åland patch network were collected during the fall survey from 4,862 nests from 449 habitat patches. These larvae were then genotyped for 45 SNPs among which 40 were putatively neutral markers (Fountain et al. 2018; DiLeo et al. 2018) and were used thereafter for our genetic analyses. To select our focal patches, we chose patches with at least 10 individuals genotyped, to ensure that the measured genetic diversity was consistent and not a result of genetic drift due to recent colonization events. We randomly sampled across nests a set of 10 genotyped individuals in focal patches to measure genetic diversity from more than one family and to avoid sample size effects. We considered the two years separately, even in cases when the same patch was sampled in both years.

Four genetic diversity indices were calculated at every focal patch using the 40 SNPs: the observed heterozygosity *Ho*, the expected heterozygosity *He*, the inbreeding coefficient *F*_*IS*_, and patch-specific *F*_*ST*_ (i.e. population-specific *F*_*ST*_, as defined in Weir & Goudet 2017). The heterozygosity coefficients provide a direct measurement of genetic diversity at the patch level, while the inbreeding coefficient and *F*_*ST*_ give information about the genetic consequences of the population dynamics and the differentiation of the subpopulations (Weir & Goudet 2017; Kitada et al. 2021). All analysis and data filtering, as well as the graphs and results presentation, were performed in R (R Core Team, 2021) using the RStudio software (RStudio Team, 2020). The genetic indices were calculated with the *Hierfstat* package (Goudet 2005).

### Definition of the study scale

We first determined the appropriate landscape scale in which the effect of habitat amount and fragmentation on genetic diversity should be assessed, as proposed by Jackson & Fahrig (2015) and Martin et al. (2021). We estimated the correlation between the four genetic diversity indices measured at the focal patches and the number of suitable fragments in the surrounding landscapes. For this, circular landscapes with radii ranging from 0.5 to 5 km by steps of 0.5 km were defined around the centroid of each focal patch (Fig. 2). For each of these landscape radii, we calculated both the total amount of suitable habitat (i.e., the total area of habitat contained in the habitat fragments) and the total number of fragments. We then performed simple linear regressions using the number of fragments as the independent variable, and one of the genetic diversity indices as the response variable for each radius. We assessed the proportion of variation explained by each model r-squares (*r*^*2*^) obtained from these regressions at all the different radii and for each of the genetic diversity indices. As suggested by Jackson & Fahrig (2015) and Martin et al. (2021), the radius for which the *r*^*2*^ was the highest was selected as our final scale of effect and used to define the local landscape surrounding each focal patch. The linear models were performed with *lme4* (Bates et al. 2022) and the R *stats* package. The calculations of the number of fragments and total habitat amount in the surrounding landscape of the focal patches at different scales were calculated using ArcGIS v. 10.8 (ESRI, Redlands, CA, USA, 2019) and QGIS v. 3.20 (QGIS, Open Source Geospatial Foundation Project, Chicago, USA, 2021).

**Figure 2.**
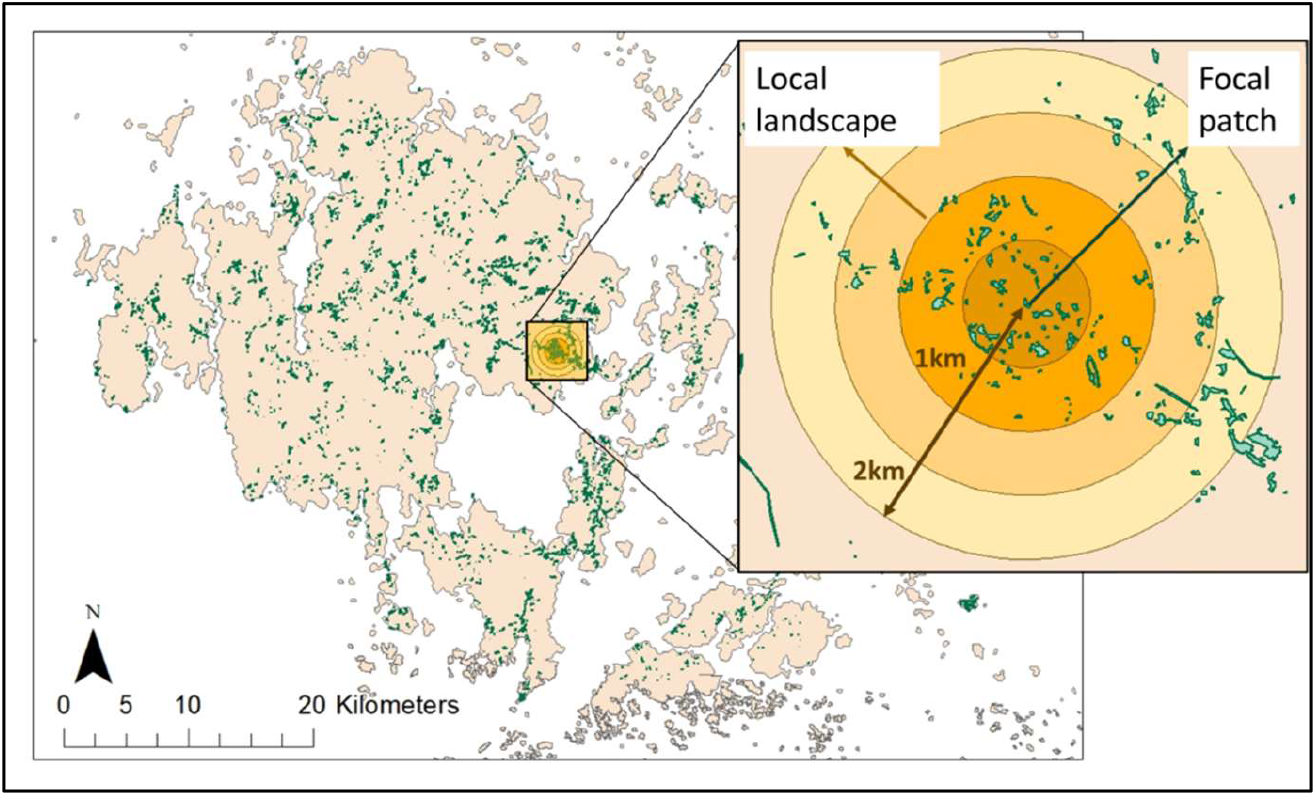
Estimation of the spatial scale with the strongest relationship between habitat amount and the genetic diversity of the Glanville fritillary butterfly in the Åland islands. Around the centroid of the focal patch, circular landscapes of radius ranging from 0.5 to 5 km by steps of 0.5 km were defined. Each green polygon corresponds to a habitat fragment of the Glanville fritillary butterfly and the white surroundings has been characterized as non-habitat.

### Testing the effects of habitat amount and fragmentation *per se* on genetic diversity

We first tested for a potential correlation between genetic diversity, habitat amount, and fragmentation at the landscape level with linear mixed-effects models. We built a model separately for each of the genetic diversity indices calculated for the focal patches as the response variable. In these models, we included the total habitat amount within the local landscape, the number of fragments within the local landscape, and their interaction as a covariate. We additionally included the focal patch size as an explanatory variable, as a large number of previous studies have shown that patch size predicts occupancy, extinction, and colonization in this system (Hanski et al. 1994; Thornton et al. 2011; Schulz et al. 2020) but few studies have taken a local-landscape approach; it is therefore unknown how important patch size is relative to landscape-scale habitat amount. Finally, the year was included as a fixed factor and the focal patch as a random effect.

As our aim was to dissociate the effects of fragmentation *per se* from the total habitat amount, we followed the Martin et al. (2021) protocol to deal with the correlation between these two variables. First, we selected a subset of local landscapes from the entire dataset in which the correlation between the total habitat amount and the number of fragments was low. As our data presented a bimodal distribution for the total habitat amount, two different subsets were defined: one including local landscapes with a small total habitat amount (0.135-0.214 Km^2^), and another with a large total habitat amount (0.449-0.521 Km^2^) in the local landscapes (Supplementary Material Fig. S1). The effect of fragmentation *per se* on the genetic diversity indices was tested separately for these two sub-datasets. For each subset, we performed, similarly as for the whole dataset, four linear mixed-effects models with each of the genetic diversity indices as the response variable. We included as explanatory variables the focal patch size, the habitat amount, the number of fragments and their interaction; as a fixed effect, the year; and as a random effect, the patch. We checked that neither of the response variables presented spatial autocorrelation.

## Results

In total, 261 out of the 449 patches had 10 or more individuals genotyped, for which we computed the genetic indices. The final dataset included 189 different patches as some were occupied during both years.

The analysis of the scale of effect showed the spatial scale at which the four genetic diversity indices present a maximum correlation with the number of fragments. The correlation between the number of fragments and *F*_*ST*_ or *He* was highest at a 3.5 km radius (*r*^*2*^ = 0.073 and *r*^*2*^ = 0.069 respectively; Suppl. Mat. Fig. S2). The highest correlation for *F*_*IS*_ was observed at 2 km (*r*^*2*^ = 0.029), and at 2.5 km for *Ho* (*r*^*2*^ = 0.069). Taking a more conservative approach, we thus chose a 3.5 km radius as our study scale for testing the habitat amount hypothesis.

Within our landscapes of 3.5 km radius, the number of fragments ranged from 30 to 291, and the total habitat amount from 0.053 to 0.613 km^2^. We first looked at the influence of our variables, including habitat amount and fragmentation *per se*, on the four genetic indices using the whole dataset. The sampling year did not affect most of the genetic diversity estimates, only *He*, with a relatively small coefficient (0.006, SE = 0.003, *p* = 0.036) (Suppl. Mat. Table S1). Using AIC criteria, we fitted a second set of models to select the best model by removing the non-significant variables. The focal patch size showed a significant positive effect on *He* (0.612, SE = 0.257, *p* = 0.018) and a negative effect on *F*_*ST*_ (−1.527, SE = 0.648, *p* = 0.020; Table 1). The habitat amount within the local landscape had a positive effect on *He* (0.046, SE = 0.013, *p* < 0.001), *Ho* (0.0517, SE = 0.015, *p* < 0.001), and *F*_*IS*_ (0.299, SE = 0.113, *p* = 0.009) while it had a negative effect on *F*_*ST*_ (−0.114, SE = 0.031, *p* < 0.001). In comparison, the number of fragments had a weak association with genetic indices. Increasing number of fragments tended to increase *F*_*ST*_ and decrease *He* and *F*_*IS*_ at higher habitat amounts, however main effects of the number of fragments and its interaction with the total habitat amount were non-significant in models (Fig. 3).

**Table 1.**
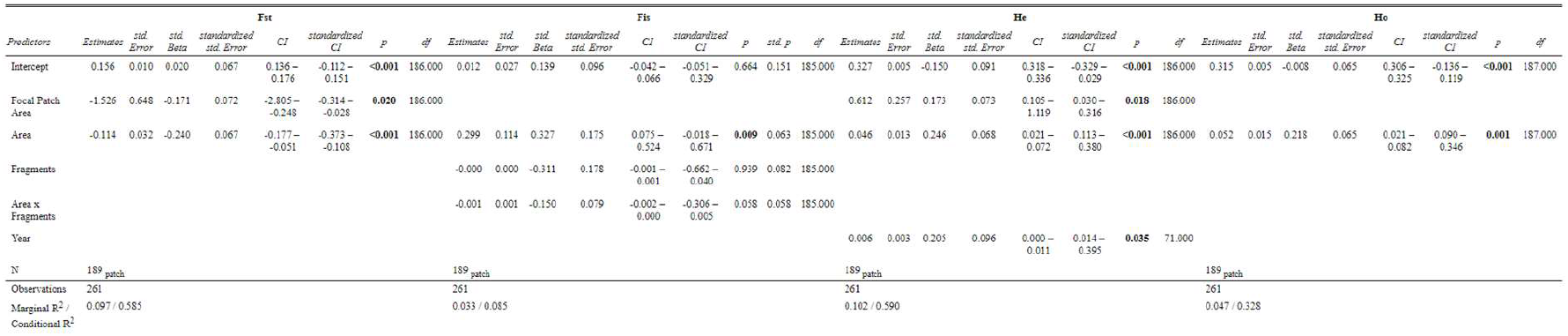
Results of the fitted linear mixed-effects models relating each of the four genetic diversity indices (*F*_*IS*_, *F*_*ST*_, *He* and *Ho*) in the Glanville fritillary butterfly to year of larval collection, focal patch size, habitat amount, and number of fragments within a landscape of 3.5 km radius.

**Figure 3.**
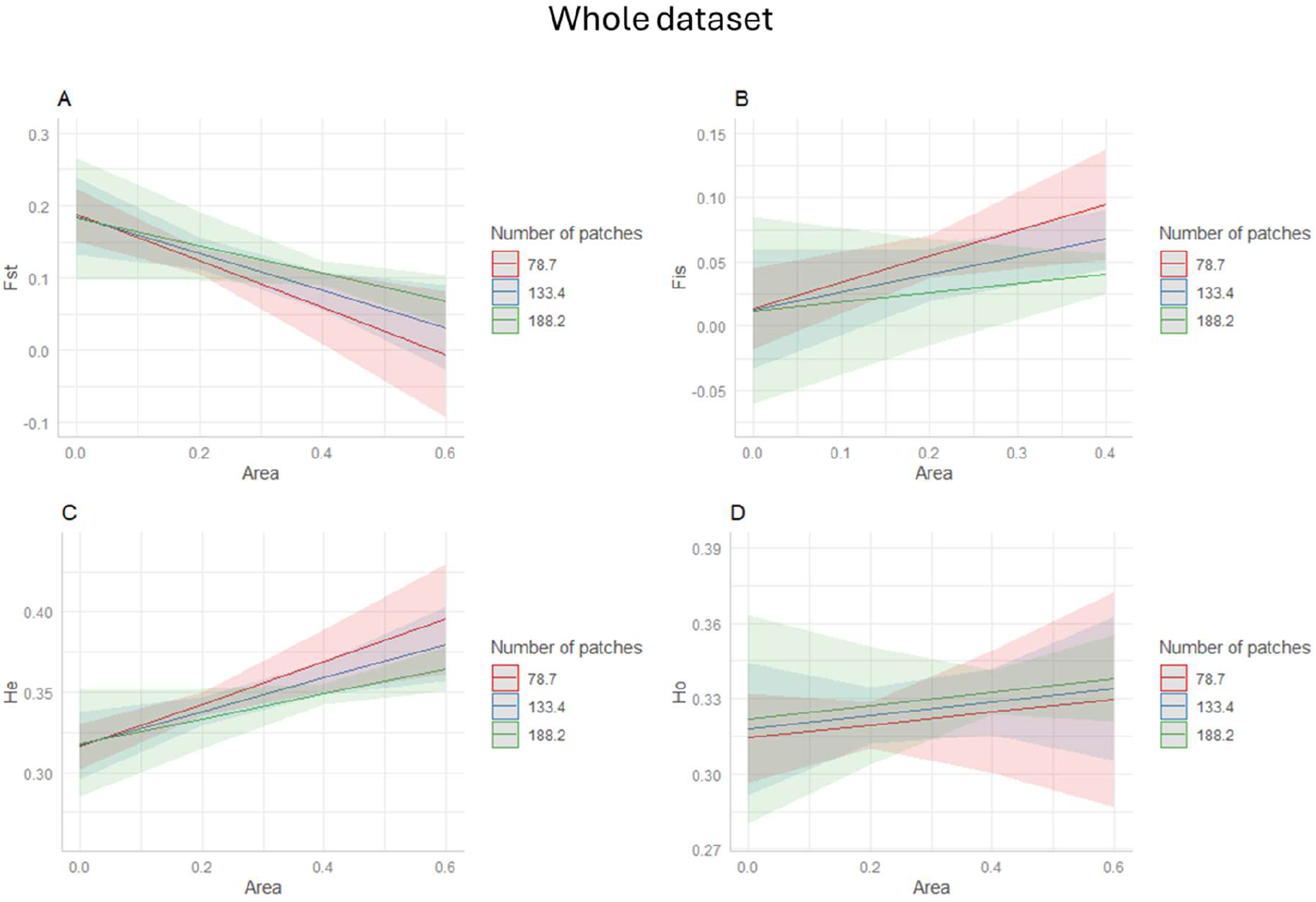
Predicted responses of the Glanville fritillary butterfly’s genetic diversity (A) *F*_*ST*_; (B) *F*_*IS*_; (C) *He*; and (D) *Ho* of focal patches to habitat amount and fragmentation *per se* (three groups of hypothetical number of patches), according to the models developed with the whole dataset (N=261).

The number of fragments and the total habitat amount within the local landscape were highly correlated (*r*^*2*^ = 0.873, *p* < 0.001; Suppl. Mat. Fig. S3). The correlation between the two variables was reduced to 0.258 for the small total habitat amount data subset, and -0.026 for the large total habitat amount, with each subset containing 97 and 63 focal patches, respectively. We repeated the same models using this time each of the two subsets. We found no significant effect of either habitat amount or fragmentation *per se* on the genetic diversity indices with either of the subsets (see Suppl. Mat. Tables S2 and S3). In the models performed with the small total habitat amount subset, there was a tendency for habitat amount to reduce *F*_*ST*_ and increase *Ho* and *He* (Fig. 4). The tendencies for the effect of fragmentation *per se* were generally weaker. Higher fragmentation tended to be associated with lower *F*_*IS*_ and higher *He* at lower values of habitat amount for the small total habitat amount subset (Fig. 4), and with lower *F*_*ST*_ and higher *Ho* for the large total habitat amount subset (Fig. S4).

**Figure 4.**
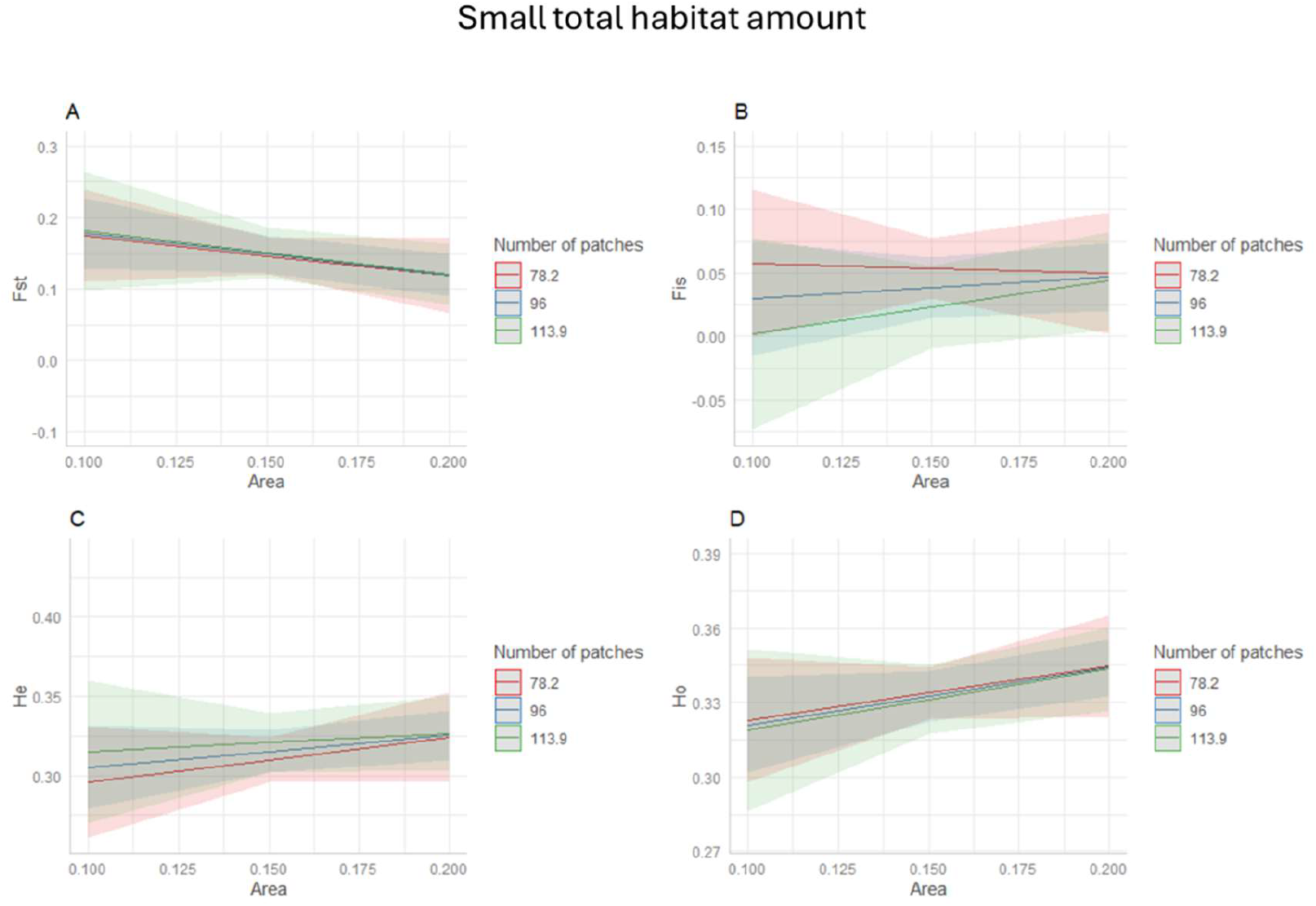
Predicted responses of the Glanville fritillary butterfly’s genetic diversity (A) *F*_*ST*_; (B) *F*_*IS*_; (C) *He*; and (D) *Ho* of focal patches to habitat amount and fragmentation *per se*, from the models developed with the subset of small total habitat amount within local landscapes (0.134-0.214 Km^2^, N = 98).

## Discussion

Habitat loss and fragmentation are main drivers of biodiversity loss, and they usually occur simultaneously (Huxel & Hastings 1999; Fahrig 2003; Wagner et al. 2021). Disentangling their distinct impact on species diversity is, however, important, as this will allow us to optimize the effectiveness of environmental management and conservation policies (Riva et al. 2024). To maintain biodiversity across scales, we also need to understand the distinct impacts of habitat amount and habitat fragmentation on genetic diversity, which is a pre-requisite for future adaptation, and hence, the long-term survival of natural populations. In our study, we found habitat amount to have a positive effect on genetic diversity both at the landscape and at the focal patch level. However, habitat amount and fragmentation were highly correlated, and we therefore could not discount the possibility that fragmentation impacts genetic diversity. When we subsampled landscapes to control for the correlation between habitat amount and fragmentation, habitat amount tended to have a positive effect, while fragmentation *per se* showed a negligible or even slightly positive effect on the neutral genetic diversity of the Glanville fritillary butterfly. We conclude that the number of fragments in the landscape has little or no effect on species’ genetic diversity after accounting for the total amount of habitat available.

### Landscape size and Glanville fritillary butterfly’s biology

The scale of effect is defined as the spatial extent at which the landscape structure best predicts the response of interest (Jackson & Fahrig 2015; Martin et al. 2021). It ensures that the landscape is measured at the appropriate scale (Jackson & Fahrig 2012). In the Åland study system, the scale at which the genetic diversity of the Glanville fritillary butterfly was responding the most to fragmentation *per se* was at 3.5 km radius. This is identical to what was found in a recent study in this system testing the effects of fragmentation per se on patch occupancy using 20 year of demographic data (Galán-Acedo et al. 2024). The multiscale analysis of the local landscape for the four different genetic indices, however, showed that the relationship between genetic diversity and the landscape extent varied depending on the genetic indicator considered. The scale of effect was the shortest for the inbreeding coefficient, *F*_*IS*_, while the largest scale of effect was found for the expected heterozygosity, *He*, and the genetic differentiation coefficient, *F*_*ST*_. When simulating population responses to habitat amount and fragmentation *per se*, Jackson & Fahrig (2014) also obtained different scales of effect depending on the response of interest with genetic diversity presenting the largest scale. It has been argued that the scale of effect is primarily a function of species mobility and that dispersal distance would be its strongest predictor (Jackson & Fahrig 2012), with the scale of effect being at least equal or higher than a species average distances (Jackson & Fahrig 2015). The observed scales of effect in our study are in line with the reported average dispersal distances of the Glanville fritillary butterfly (Hanski et al. 1995; Fountain et al. 2018; DiLeo et al. 2018). The estimated average dispersal distance for the adults is 1.5 km (DiLeo et al. 2018, 2022), with the highest breeding dispersal distances reaching even 10 km based on genetic assignment tests (DiLeo et al. 2022). This suggests that we are capturing the appropriate landscape size to disentangle the role of fragmentation *per se* from habitat amount.

### Effects of habitat amount and fragmentation *per se* on genetic diversity

As suggested in the habitat amount hypothesis (Fahrig 2013) and metapopulation persistence theory (With 2004), the total habitat amount within the landscape seemed to be the main factor affecting the genetic diversity of the Glanville fritillary butterfly. Jackson & Fahrig (2014) found a similar trend in genetic diversity in their theoretical study. Our results suggest that with increasing habitat amount, expected and observed heterozygosity as well as inbreeding increased, while population differentiation decreased. After accounting for the correlation between habitat amount and the number of fragments within local landscapes (Martin et al. 2021), the association between habitat amount and genetic diversity was less clear. Within local landscapes with small total habitat amount, the genetic diversity of the Glanville fritillary butterfly had the tendency to respond positively to habitat amount, independently from fragmentation *per se*. An increase in the total habitat amount in the local landscape can thus reduce the rate of genetic drift, improve gene flow between patches (Keyghobadi et al. 2005; Lebigre et al. 2022) and in turn, favor the evolutionary potential of the local population (Cheptou et al. 2017). However, we did not observe a trend indicating a positive effect of habitat amount within local landscapes with a large habitat amount (Suppl. Mat. Fig. S3). This could be because the differences in habitat amount can present more noticeable effects when the habitat is already scarce (Hanski 2011a). Thus, if the amount of habitat in the local landscape is above a certain threshold, it is possible that the effect of small differences in habitat amount is not big enough to be detected. However, as the sample size of the large habitat amount subset was much smaller, we cannot exclude a low statistical power to observe any effects of habitat amount.

The effects of fragmentation *per se* on genetic diversity of the Glanville fritillary butterfly were generally weak and non-significant. However, we acknowledge that due to a very high correlation between habitat amount and fragmentation, standard errors of estimated effects of fragmentation from the model using the complete dataset are likely inflated and unstable and thus must be interpreted with caution (Dormann et al. 2013). After controlling for the correlation with habitat amount, we found no clear effect of fragmentation *per se* on genetic diversity, despite our attempt to maintain as much variability in the number of fragments as possible and to reduce the variability in the total habitat amount. Instead, more often than not, we observed patterns consistent with a neutral or even positive effect of fragmentation on genetic diversity. The genetic diversity of species with a fragmented distribution, including the Glanville Fritillary population in the Åland islands, depends greatly on the rate of migration between habitat patches. Sufficient gene flow between habitat patches can reduce genetic drift and thus prevent the negative effects of fragmentation (Amos & Harwood 1998). In this particular study system, fragmentation *per se* has also been found to have positive effects on patch occupancy, potentially attributed to a decrease in the mean distances between patches in landscapes with more patches (Galán-Acedo et al. 2024) (i.e., when fragmentation is defined as the number of habitat fragments, and for a constant amount of habitat, an increase in fragmentation implies that the mean separation between patches is reduced (Fahrig 2017)). Thus, landscapes with more habitat patches could present higher overall connectivity, leading to higher migration rates and, consequently, higher genetic diversity.

Finally, in line with previous ecological observations on the role of increased habitat patch area in the Glanville fritillary butterfly (Hanski 1999), we found that the size of the focal patch was affecting some of the genetic diversity indices, namely *F*_*ST*_ and *He*. When the size of the focal patch increased, patch genetic differentiation decreased while expected heterozygosity increased, indicating that more immigrants from the surrounding patches are colonizing larger focal patches. Larger and well-connected habitat patches tend to support higher occupancy and population abundance (Schulz et al. 2020). On the other hand, small habitat fragment patches do not often have a very high occupancy rate, high population abundance nor do they stay occupied for many consecutive years (Dallas et al. 2020), and instead are more likely to work as stepping stone patches. Smaller focal patches will likely contain smaller subpopulations, show a lower immigration rate, and increase genetic drift which can lead to genetically more differentiated subpopulations.

### Disentangling habitat amount from fragmentation *per se* effects on genetic diversity of wild populations

Characteristics and quality of local habitat patches may be important determinants of effective population size (*Ne*) and consequently, genetic diversity (Crawford & Keyghobadi 2018). Studies have shown the importance of patch quality and patch connectivity in butterflies on population density (Turlure et al. 2010; Crawford & Keyghobadi 2018), successful dispersal within the landscape (With 2004; DiLeo et al. 2022), patch occupancy (Dennis & Eales 1997; Hanski et al. 2017; Galán-Acedo et al. 2024) as well as genetic diversity (Lebigre et al. 2022). When only taking into account the habitat amount available within the local landscape, patch quality is not considered, while patch connectivity is assumed to increase with the total habitat amount but is not directly measured (Fahrig 2013). For small organisms like insects, direct measures of patch connectivity (e.g. using the incidence function model) have been found to be good estimators of both dispersibility and maintenance in the landscape while patch quality, an estimator of individual settlement (DiLeo et al. 2022). If individuals cannot colonize some patches within the landscape due to their isolation or poor quality, it will decrease the effective population size the landscape can hold, potentially exposing the population to a higher risk of extinction. In turn, the population is likely to be more sensitive to stochasticity and random events which could lead to a reduce genetic diversity. This might become even more important as extreme events (i.e. droughts, flooding) are increasing (Calvin et al. 2023), directly affecting insects (Wagner et al. 2021) including the Glanville fritillary butterfly (Bergen et al. 2020). Measuring the genetic diversity of a species at the landscape level might then give better insight into the overall viability of the population by providing more representative information about the maintenance of genetic diversity while the patch scale approach would provide information on species movement, barriers (Holderegger & Wagner 2008) and density fluctuations between patches. Genetic studies combining patch-based and landscape-based approaches will thus permit more accurate predictions of the responses of a species’ genetic diversity to landscape characteristics. Such studies will be especially relevant as Riva & Fahrig (2022) found opposite patterns in species diversity when using either a landscape-scale or patch-scale approach.

Finally, we lost substantial statistical power by subsetting our data to control for high correlations between habitat amount and fragmentation. Future studies could avoid this limitation by selecting *a priori* and sampling in landscapes with more contrasting characteristics (i.e., including more landscapes with high habitat amount and low fragmentation, and landscapes with low habitat amount and high fragmentation). Considering the urgency to protect the remaining biodiversity (Cowie et al. 2022), such an approach would allow us to determine if measuring only the effect of habitat amount is a more efficient method to develop appropriate conservation measures than trying to consider the full complexity of the landscape (Seibold et al. 2017).

## Conclusion

Studies testing the effect of fragmentation *per se* on species’ genetic diversity using wild populations are still scarce (but see Connor et al. 2022), and to our knowledge, do not exist for insects. Our results suggest that the total habitat amount available within the landscape affects genetic diversity positively, while fragmentation *per se* seems to have a neutral effect on genetic diversity. These outcomes thus provide support for the benefit of first prioritizing the conservation of large enough habitat availability within the local landscape and in a second step, its habitat configuration (Riva et al. 2024). Overall, our study demonstrates the complexity of assessing the impacts of fragmentation *per se* on genetic diversity estimates, even in a species for which extensive landscape and genetic datasets are available. Future studies testing the habitat amount hypothesis at both neutral and functional genetic diversity in more contrasted landscapes would give more insight in the separate role of habitat amount and fragmentation *per se* on species.

