## Supplementary Material file for "Relative Effects of Habitat Amount and Fragmentation *Per Se* on the Genetic Diversity of The Glanville Fritillary Butterfly"

Figure S1. Distribution of the total habitat area (km$^2$) within each circular local landscape of 3.5 km radius. The defined data subsets for (A) the small total habitat amount (0.134-0.214 Km$^2$; N = 88), and (B) the large total habitat amount (0.449-0.521 Km$^2$; N = 56) are indicated in red.

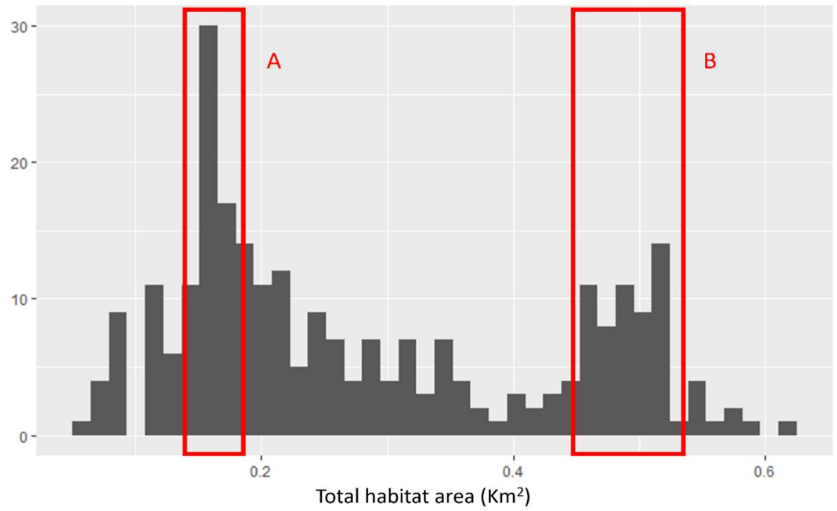


Figure S2. Estimation of the scale of effect for the Glanville fritillary in the Åland islands. The $r^2$ represents the relationship between each of the four genetic indices (*Ho*, *He*, $F_{IS}$ and $F_{ST}$) and the number of fragments within a defined local landscape radius.

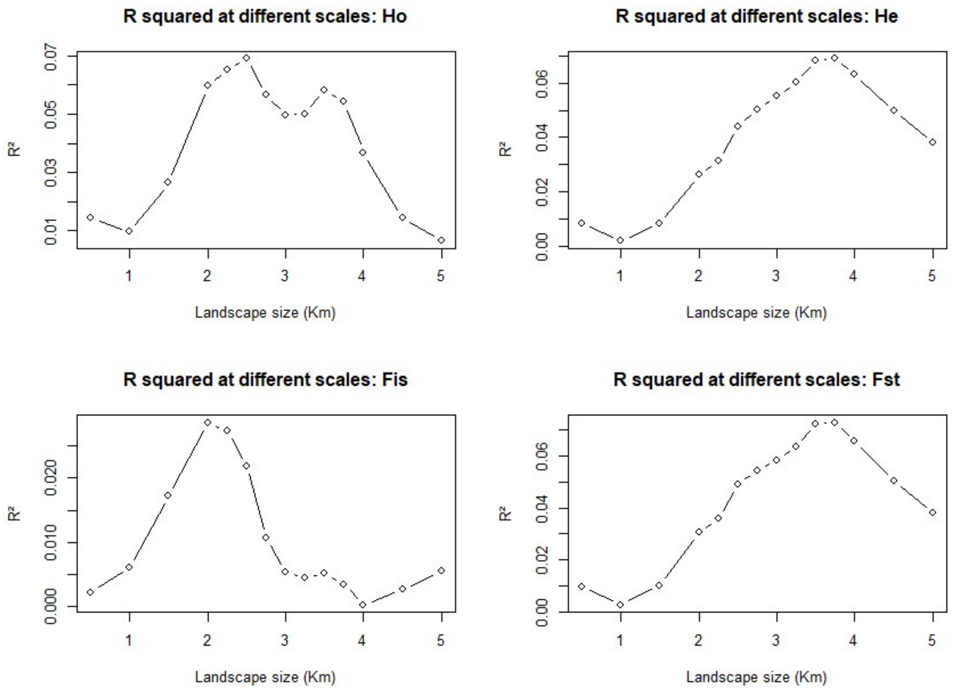

Figure S3. Relationship between the total habitat amount and the number of fragments within local landscapes of 3.5 km radius around the focal patch for which the genetic diversity indices were calculated ($r^2 = 0.87$).

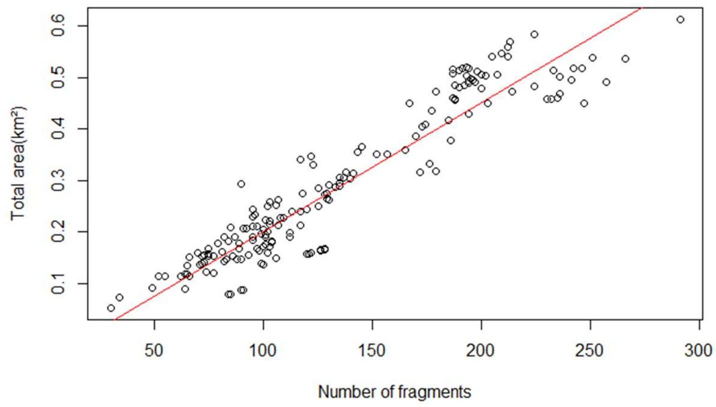


Figure S4. Predicted responses of the Glanville fritillary butterfly's genetic diversity (A) $F_{ST}$; (B) $F_{IS}$; (C) *He*; and (D) *Ho* of focal patches to habitat amount and fragmentation *per se,* from the subset of large total habitat amount within local landscapes (0.449-0.521 $Km^2$; N = 57).

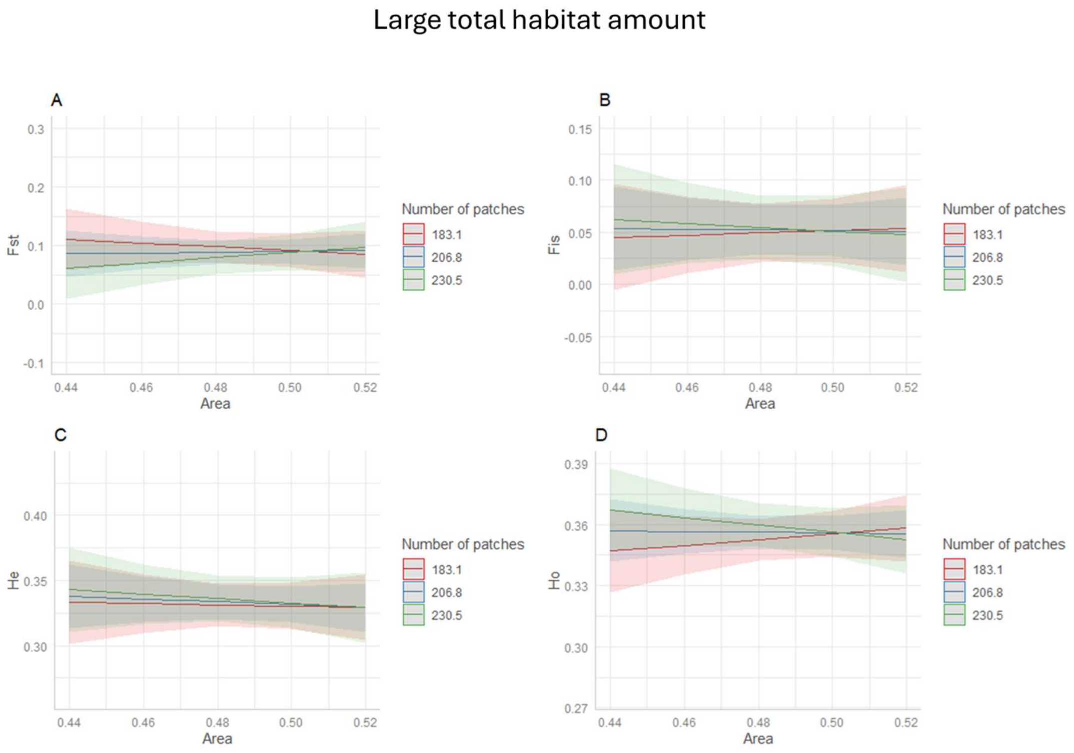

Table S1. Results of the linear mixed-effects models relating each of the four genetic diversity indices ($F_{IS}$, $F_{ST}$, *He* and *Ho*) to the year of larval collection, focal patch size, habitat amount, and number of fragments within a local landscape of 3.5 km radius. The models include significant and non-significant variables.

**Fst**

| *Predictors* | *Estimates* | *std. Error* | *std. Beta* | *standardized std. Error* | *CI* | *standardized CI* | *p* | *std. p* | *df* |
|---|---|---|---|---|---|---|---|---|---|
| Intercept | 0.199 | 0.032 | -0.027 | 0.119 | 0.136 – 0.261 | -0.261 – 0.207 | **<0.001** | 0.821 | 184.000 |
| Focal Patch Area | -1.367 | 0.651 | -0.153 | 0.073 | -2.650 – -0.083 | -0.296 – -0.009 | **0.037** | **0.037** | 184.000 |
| Area | -0.418 | 0.134 | -0.539 | 0.190 | -0.682 – -0.153 | -0.914 – -0.163 | **0.002** | **0.005** | 184.000 |
| Fragments | -0.000 | 0.000 | 0.233 | 0.191 | -0.001 – 0.001 | -0.143 – 0.609 | 0.892 | 0.224 | 184.000 |
| Area x Fragments | -0.009 | 0.007 | -0.128 | 0.095 | -0.022 – 0.004 | -0.317 – 0.062 | 0.183 | 0.183 | 71.000 |
| Year | 0.001 | 0.001 | 0.139 | 0.081 | -0.000 – 0.003 | -0.021 – 0.299 | 0.089 | 0.089 | 184.000 |
| N | 189 patch | | | | | | | | |
| Observations | 261 | | | | | | | | |
| Marginal $R^2$ / Conditional $R^2$ | 0.123 / 0.598 | | | | | | | | |

**Fis**

| *Predictors* | *Estimates* | *std. Error* | *std. Beta* | *standardized std. Error* | *CI* | *standardized CI* | *p* | *std. p* | *df* |
|---|---|---|---|---|---|---|---|---|---|
| Intercept | 0.015 | 0.028 | 0.186 | 0.126 | -0.040 – 0.070 | -0.062 – 0.434 | 0.598 | 0.141 | 184.000 |
| Focal Patch Area | 0.001 | 0.524 | 0.000 | 0.065 | -1.032 – 1.035 | -0.127 – 0.128 | 0.998 | 0.998 | 184.000 |
| Area | 0.297 | 0.116 | 0.318 | 0.179 | 0.069 – 0.526 | -0.036 – 0.672 | **0.011** | 0.078 | 184.000 |
| Fragments | -0.000 | 0.000 | -0.306 | 0.181 | -0.001 – 0.001 | -0.663 – 0.050 | 0.962 | 0.091 | 184.000 |
| Area x Fragments | -0.005 | 0.008 | -0.074 | 0.126 | -0.021 – 0.011 | -0.324 – 0.177 | 0.559 | 0.559 | 71.000 |
| Year | -0.001 | 0.001 | -0.152 | 0.079 | -0.002 – 0.000 | -0.309 – 0.005 | 0.057 | 0.057 | 184.000 |
| N | 189 patch | | | | | | | | |
| Observations | 261 | | | | | | | | |
| Marginal $R^2$ / Conditional $R^2$ | 0.034 / 0.087 | | | | | | | | |

**He**

| *Predictors* | *Estimates* | *std. Error* | *std. Beta* | *standardized std. Error* | *CI* | *standardized CI* | *p* | *std. p* | *df* |
|---|---|---|---|---|---|---|---|---|---|
| Intercept | 0.312 | 0.012 | -0.013 | 0.119 | 0.288 – 0.337 | -0.248 – 0.221 | **<0.001** | 0.910 | 184.000 |
| Focal Patch Area | 0.532 | 0.257 | 0.150 | 0.073 | 0.025 – 1.039 | 0.007 – 0.293 | **0.040** | **0.040** | 184.000 |
| Area | 0.172 | 0.053 | 0.556 | 0.190 | 0.067 – 0.276 | 0.181 – 0.930 | **0.001** | **0.004** | 184.000 |
| Fragments | 0.000 | 0.000 | -0.250 | 0.190 | -0.000 – 0.000 | -0.626 – 0.125 | 0.902 | 0.190 | 184.000 |
| Area x Fragments | 0.006 | 0.003 | 0.204 | 0.095 | 0.000 – 0.011 | 0.014 – 0.395 | **0.036** | **0.036** | 71.000 |
| Year | -0.001 | 0.000 | -0.146 | 0.081 | -0.001 – 0.000 | -0.306 – 0.014 | 0.073 | 0.073 | 184.000 |
| N | 189 patch | | | | | | | | |
| Observations | 261 | | | | | | | | |
| Marginal $R^2$ / Conditional $R^2$ | 0.126 / 0.592 | | | | | | | | |

**Ho**

| *Predictors* | *Estimates* | *std. Error* | *std. Beta* | *standardized std. Error* | *CI* | *standardized CI* | *p* | *std. p* | *df* |
|---|---|---|---|---|---|---|---|---|---|
| Intercept | 0.306 | 0.016 | -0.134 | 0.123 | 0.275 – 0.337 | -0.377 – 0.110 | **<0.001** | 0.281 | 184.000 |
| Focal Patch Area | 0.473 | 0.310 | 0.106 | 0.069 | -0.139 – 1.085 | -0.031 – 0.243 | 0.129 | 0.129 | 184.000 |
| Area | 0.025 | 0.066 | 0.111 | 0.187 | -0.105 – 0.155 | -0.257 – 0.479 | 0.705 | 0.553 | 184.000 |
| Fragments | 0.000 | 0.000 | 0.112 | 0.187 | -0.000 – 0.000 | -0.257 – 0.482 | 0.684 | 0.549 | 184.000 |
| Area x Fragments | 0.007 | 0.004 | 0.200 | 0.112 | -0.001 – 0.015 | -0.024 – 0.423 | 0.079 | 0.079 | 71.000 |
| Year | 0.000 | 0.000 | 0.002 | 0.081 | -0.001 – 0.001 | -0.157 – 0.162 | 0.978 | 0.978 | 184.000 |
| N | 189 patch | | | | | | | | |
| Observations | 261 | | | | | | | | |
| Marginal $R^2$ / Conditional $R^2$ | 0.068 / 0.356 | | | | | | | | |

Table S2. Results of the linear mixed-effects models performed with the small total amount of habitat subset, relating each of the four genetic diversity indices ($F_{IS}$, $F_{ST}$, *He* and *Ho*) to the year of larval collection, focal patch size, habitat amount, and number of fragments within a local landscape of 3.5 km radius. The models include significant and non-significant variables.

**Fst**

| *Predictors* | *Estimates* | *std. Error* | *std. Beta* | *standardized std. Error* | *CI* | *standardized CI* | *p* | *std. p* |
|---|---|---|---|---|---|---|---|---|
| Intercept | 0.211 | 0.390 | 0.175 | 0.177 | -0.568 – 0.991 | -0.179 – 0.530 | 0.590 | 0.326 |
| Fragments | 0.000 | 0.004 | 0.027 | 0.137 | -0.008 – 0.009 | -0.247 – 0.300 | 0.940 | 0.846 |
| Area | -0.459 | 2.446 | -0.207 | 0.113 | -5.344 – 4.425 | -0.433 – 0.018 | 0.852 | 0.071 |
| Area x Fragments | -1.302 | 1.417 | -0.104 | 0.114 | -4.132 – 1.528 | -0.331 – 0.122 | 0.362 | 0.362 |
| Focal Patch Area | -0.015 | 0.013 | -0.230 | 0.196 | -0.041 – 0.011 | -0.632 – 0.171 | 0.250 | 0.250 |
| Year | -0.001 | 0.025 | -0.008 | 0.163 | -0.052 – 0.050 | -0.333 – 0.317 | 0.961 | 0.961 |
| N | 70 patch | | | | | | | |
| Observations | 98 | | | | | | | |
| Marginal $R^2$ / Conditional $R^2$ | 0.060 / 0.311 | | | | | | | |

**Fis**

| *Predictors* | *Estimates* | *std. Error* | *std. Beta* | *standardized std. Error* | *CI* | *standardized CI* | *p* | *std. p* |
|---|---|---|---|---|---|---|---|---|
| Intercept | 0.297 | 0.350 | 0.029 | 0.179 | -0.403 – 0.997 | -0.328 – 0.386 | 0.400 | 0.872 |
| Fragments | -0.003 | 0.004 | -0.174 | 0.135 | -0.010 – 0.004 | -0.443 – 0.094 | 0.428 | 0.200 |
| Area | -1.159 | 2.196 | 0.067 | 0.111 | -5.546 – 3.227 | -0.155 – 0.289 | 0.599 | 0.549 |
| Area x Fragments | -0.276 | 1.271 | -0.024 | 0.111 | -2.814 – 2.261 | -0.246 – 0.198 | 0.829 | 0.829 |
| Focal Patch Area | -0.005 | 0.012 | -0.084 | 0.202 | -0.029 – 0.020 | -0.497 – 0.330 | 0.681 | 0.681 |
| Year | 0.014 | 0.023 | 0.097 | 0.159 | -0.032 – 0.060 | -0.222 – 0.415 | 0.547 | 0.547 |
| N | 70 patch | | | | | | | |
| Observations | 98 | | | | | | | |
| Marginal $R^2$ / Conditional $R^2$ | 0.049 / 0.238 | | | | | | | |

**Ho**

| *Predictors* | *Estimates* | *std. Error* | *std. Beta* | *standardized std. Error* | *CI* | *standardized CI* | *p* | *std. p* |
|---|---|---|---|---|---|---|---|---|
| Intercept | 0.187 | 0.213 | -0.203 | 0.173 | -0.239 – 0.613 | -0.548 – 0.142 | 0.383 | 0.245 |
| Fragments | 0.001 | 0.002 | 0.108 | 0.140 | -0.003 – 0.005 | -0.172 – 0.388 | 0.660 | 0.444 |
| Area | 0.640 | 1.336 | 0.134 | 0.116 | -2.028 – 3.307 | -0.097 – 0.364 | 0.634 | 0.252 |
| Area x Fragments | 0.667 | 0.776 | 0.101 | 0.117 | -0.884 – 2.218 | -0.133 – 0.334 | 0.394 | 0.394 |
| Focal Patch Area | 0.010 | 0.006 | 0.302 | 0.182 | -0.002 – 0.023 | -0.071 – 0.676 | 0.109 | 0.109 |
| Year | -0.005 | 0.014 | -0.055 | 0.167 | -0.032 – 0.023 | -0.389 – 0.278 | 0.742 | 0.742 |
| N | 70 patch | | | | | | | |
| Observations | 98 | | | | | | | |
| Marginal $R^2$ / Conditional $R^2$ | 0.074 / 0.439 | | | | | | | |

**He**

| *Predictors* | *Estimates* | *std. Error* | *std. Beta* | *standardized std. Error* | *CI* | *standardized CI* | *p* | *std. p* |
|---|---|---|---|---|---|---|---|---|
| Intercept | 0.314 | 0.153 | -0.225 | 0.177 | 0.008 – 0.619 | -0.579 – 0.128 | **0.044** | 0.208 |
| Fragments | -0.000 | 0.002 | -0.039 | 0.136 | -0.003 – 0.003 | -0.310 – 0.232 | 0.908 | 0.776 |
| Area | 0.157 | 0.958 | 0.209 | 0.112 | -1.756 – 2.070 | -0.015 – 0.432 | 0.870 | 0.067 |
| Area x Fragments | 0.492 | 0.555 | 0.100 | 0.112 | -0.616 – 1.601 | -0.125 – 0.324 | 0.378 | 0.378 |
| Focal Patch Area | 0.008 | 0.005 | 0.307 | 0.196 | -0.002 – 0.018 | -0.096 – 0.709 | 0.130 | 0.130 |
| Year | 0.001 | 0.010 | 0.012 | 0.161 | -0.019 – 0.021 | -0.309 – 0.334 | 0.939 | 0.939 |
| N | 70 patch | | | | | | | |
| Observations | 98 | | | | | | | |
| Marginal $R^2$ / Conditional $R^2$ | 0.068 / 0.301 | | | | | | | |

Table S3. Results of the linear mixed-effects models performed with the large total amount of habitat subset, relating each of the four genetic diversity indices ($F_{IS}$, $F_{ST}$, *He* and *Ho*) to the year of larval collection, focal patch size, habitat amount, and number of fragments within a local landscape of 3.5 km radius. The models include significant and non-significant variables.

| Predictors | Fst Estimates | std. Error | std. Beta | standardized std. Error | CI | standardized CI | p | std. p |
|---|---|---|---|---|---|---|---|---|
| Intercept | 1.780 | 1.470 | -0.009 | 0.180 | -1.208 – 4.768 | -0.375 – 0.357 | 0.234 | 0.960 |
| Fragments | -0.008 | 0.007 | -0.119 | 0.162 | -0.023 – 0.006 | -0.448 – 0.211 | 0.249 | 0.470 |
| Area | -3.330 | 3.031 | 0.028 | 0.157 | -9.490 – 2.830 | -0.290 – 0.346 | 0.280 | 0.859 |
| Area x Fragments | -2.369 | 1.552 | -0.246 | 0.161 | -5.523 – 0.784 | -0.573 – 0.081 | 0.136 | 0.136 |
| Focal Patch Area | 0.002 | 0.010 | 0.036 | 0.190 | -0.020 – 0.024 | -0.365 – 0.438 | 0.851 | 0.851 |
| Year | 0.016 | 0.015 | 0.169 | 0.149 | -0.013 – 0.046 | -0.135 – 0.472 | 0.267 | 0.267 |
| N | 39 patch | | | | | | | |
| Observations | 57 | | | | | | | |
| Marginal $R^2$ / Conditional $R^2$ | 0.091 / 0.637 | | | | | | | |

| Predictors | Fis Estimates | std. Error | std. Beta | standardized std. Error | CI | standardized CI | p | std. p |
|---|---|---|---|---|---|---|---|---|
| Intercept | -0.574 | 1.415 | 0.203 | 0.194 | -3.449 – 2.301 | -0.190 – 0.597 | 0.687 | 0.301 |
| Fragments | 0.003 | 0.007 | 0.026 | 0.143 | -0.011 – 0.017 | -0.265 – 0.317 | 0.653 | 0.856 |
| Area | 1.247 | 2.923 | -0.015 | 0.137 | -4.693 – 7.187 | -0.292 – 0.263 | 0.672 | 0.914 |
| Area x Fragments | 0.915 | 1.506 | 0.085 | 0.140 | -2.145 – 3.975 | -0.200 – 0.371 | 0.547 | 0.547 |
| Focal Patch Area | -0.025 | 0.017 | -0.403 | 0.273 | -0.060 – 0.011 | -0.978 – 0.173 | 0.158 | 0.158 |
| Year | -0.006 | 0.014 | -0.057 | 0.130 | -0.035 – 0.022 | -0.322 – 0.207 | 0.662 | 0.662 |
| N | 39 patch | | | | | | | |
| Observations | 57 | | | | | | | |
| Marginal $R^2$ / Conditional $R^2$ | 0.048 / 0.048 | | | | | | | |

| Predictors | Ho Estimates | std. Error | std. Beta | standardized std. Error | CI | standardized CI | p | std. p |
|---|---|---|---|---|---|---|---|---|
| Intercept | 0.100 | 0.891 | -0.200 | 0.190 | -1.712 – 1.911 | -0.587 – 0.187 | 0.912 | 0.301 |
| Fragments | 0.001 | 0.004 | 0.057 | 0.160 | -0.007 – 0.010 | -0.267 – 0.381 | 0.750 | 0.723 |
| Area | 0.439 | 1.839 | -0.076 | 0.153 | -3.298 – 4.175 | -0.388 – 0.235 | 0.813 | 0.622 |
| Area x Fragments | 0.506 | 0.943 | 0.085 | 0.158 | -1.410 – 2.422 | -0.236 – 0.405 | 0.595 | 0.595 |
| Focal Patch Area | 0.013 | 0.008 | 0.380 | 0.231 | -0.004 – 0.030 | -0.108 – 0.867 | 0.119 | 0.119 |
| Year | -0.003 | 0.009 | -0.044 | 0.146 | -0.021 – 0.015 | -0.341 – 0.253 | 0.766 | 0.766 |
| N | 39 patch | | | | | | | |
| Observations | 57 | | | | | | | |
| Marginal $R^2$ / Conditional $R^2$ | 0.051 / 0.417 | | | | | | | |

| Predictors | Hs Estimates | std. Error | std. Beta | standardized std. Error | CI | standardized CI | p | std. p |
|---|---|---|---|---|---|---|---|---|
| Intercept | -0.335 | 0.572 | -0.033 | 0.181 | -1.497 – 0.828 | -0.402 – 0.335 | 0.563 | 0.855 |
| Fragments | 0.003 | 0.003 | 0.115 | 0.162 | -0.002 – 0.009 | -0.213 – 0.444 | 0.229 | 0.481 |
| Area | 1.363 | 1.180 | -0.022 | 0.156 | -1.035 – 3.761 | -0.339 – 0.296 | 0.256 | 0.890 |
| Area x Fragments | 0.936 | 0.604 | 0.249 | 0.161 | -0.292 – 2.164 | -0.078 – 0.575 | 0.131 | 0.131 |
| Focal Patch Area | 0.001 | 0.004 | 0.048 | 0.195 | -0.008 – 0.010 | -0.364 – 0.460 | 0.809 | 0.809 |
| Year | -0.007 | 0.006 | -0.176 | 0.149 | -0.018 – 0.005 | -0.479 – 0.127 | 0.246 | 0.246 |
| N | 39 patch | | | | | | | |
| Observations | 57 | | | | | | | |
| Marginal $R^2$ / Conditional $R^2$ | 0.094 / 0.617 | | | | | | | |
